# *Lens* species genome assemblies reveal evolutionarily fragile chromosomes associated with pericentric inversions

**DOI:** 10.64898/2026.09.02.749020

**Authors:** A. J. A Silvestrini, L. Ramsay, K. Koh, M. Baker, M. Mau, S. Kaur, S Sudheesh, N. Rodde, L. Pasipanodya, S. Kagale, R. Stonehouse, E. von-Wettberg, K. E Bett

**Affiliations:** Department of Plant Sciences, University of Saskatchewan, Saskatoon, SK, Canada; Global Institute for Food Security, University of Saskatchewan, Saskatoon, SK, Canada; Department of Computer Science, University of Saskatchewan, Saskatoon, SK, Canada; Omics Resource Centre, Western College of Veterinary Medicine, Saskatoon, SK, Canada; Agriculture Science and Technology, AgriBio, Centre for AgriBioscience, Department of Environment, Energy & Climate Action (DEECA), Bundoora, Victoria, Australia; Univ Toulouse, INRAE, Castanet Tolosan, France; Aquatic and Crop Resource Development, National Research Council Canada, Saskatoon, SK, Canada; Department of Agriculture and Life Sciences and Gund Institute for the Environment, University of Vermont, Burlington, United States of America

## Abstract

The genus *Lens* is comprised of seven diploid species, including the cultivated crop Lentil, and six wild relatives often used in breeding programs for beneficial allele introgression. *Lens* species are proposed to exhibit considerable levels of genome structural variation both within and between species, which has implications for plant breeding and evolutionary genomics. Plant centromeres are key regions involved in chromosomal stability, where structural changes can be linked to meiotic abnormalities, resulting in inversions, translocations, and deletions of chromosomal segments. Fabeae centromeres are known to be diverse and do not fit the centromere drive model. The origins of this centromere diversity and the evolutionary trends remain unclear, but recent studies suggest that centromere structural variation plays a role in creating new sequence combinations that contribute to the unique centromere evolution patterns observed in legumes. We sequenced, assembled, and performed gene synteny analysis in the genomes of all seven *Lens* species using long-read sequencing technologies. We also located and characterized the centromeres in these genomes through ChIP-seq experiments to better understand their role in *Lens* chromosomal stability. Synteny analysis reveals that *Lens* genomes have undergone extensive rearrangements during evolution, with multiple translocations and inversions identified between species. Specific *Lens* chromosomes appear more susceptible to rearrangements over time. These same chromosomes also exhibit higher rates of rearrangement throughout *Lens* speciation. Fragile chromosomes are associated with the occurrence of pericentric inversions, likely disrupting stability. The diversity in satellite sequences in *Lens* centromeres likely results from structural changes accumulated throughout *Lens* evolution. These findings suggest that genome and centromere structure play significant roles in generating centromere diversity within this legume genus.

## Introduction

The genus *Lens* belongs to the family Fabaceae and is comprised of seven species, of which Lentil (*Lens culinaris*) is an important cool-season legume crop (Ferguson et al., 2000). Lentil is a source of complex carbohydrates, fibres, vitamins and minerals and can provide twice as much protein as wheat or rice (Laskar et al., 2019; Yadav et al., 2007). As with many legumes, *Lens* species can associate with rhizobia that fix atmospheric nitrogen in root nodules, which allows them to thrive in low nitrogen environments and benefits the sustainable production of sequential crops (Quinn, 2009). Lentil wild relatives possess characteristics of interest for introgression into the cultivated germplasm, such as biotic and abiotic stress tolerance and the potential for biofortification, increasing its nutritional qualities (Cao et al., 2024; Kumar et al., 2018; Marshall et al., 2024; Singh et al., 2018; Tullu et al., 2011; Vargas et al., 2024). *Lens* spp. are interesting for evolutionary genomics studies due to their small number and a stable ploidy level of 2n = 2x = 7 for all seven taxa (Galasso, 2003).

Comparative genomics is a powerful tool for characterizing genetic diversity. Across *Lens* spp., broader geographic distributions are associated with increased genetic diversity and extensive genome structural variation (Guerra-García, et al., 2021). Chromosomal rearrangements are structural genomic changes that involve DNA segments longer than 1 kb (Scherer et al., 2007) and, in general, can lead to the loss, duplication, inversion or translocation of chromosomal fragments in one species relative to another (Scouarnec & Gribble, 2012), which leads to reduced fertility in crosses and speciation over time. Chromosomal rearrangements, such as inversions, can also occur in evolutionarily conserved structures, including centromeres. When an inversion spans the centromeric regions of a chromosome, it is called a pericentric inversion.

Centromeres are predominantly defined epigenetically by the presence of the histone H3 variant CENH3 (also known as CENP-A). They have a conserved function of ensuring proper chromosome segregation during cell division, which contrasts with their extremely diverse eukaryotic sequence composition. The genus *Lens* belongs to the legume tribe Fabeae, which has been shown to have exceptionally high inter- and intraspecific diversity of centromeric repeats (Robledillo et al., 2020) and centromere structure (Neumann et al., 2015).

Comparative and evolutionary centromere studies are still rare, and most studies are related to putative centromeric satellite sequence evolution and not directly associated with precise centromere determination using ChIP-seq experiments (Huang et al., 2021; Kadluczka & Grzebelus, 2021; Mandáková et al., 2020; Melters et al., 2013; Montenegro et al., 2022; Zwyrtková et al., 2020). There is no record of comparative centromere analysis on an entire genus using long-read genome assembly technologies. An across-genus analysis could help decipher centromere evolution in relation to specific evolutionary processes, such as speciation due to mating barriers or centromere evolution facing larger structural variation. Here we perform such an analysis in the *genus* Lens, finding extensive variation in centromeres across the group.

## Material and Methods

### Plant Material and DNA Extraction

All accessions that were sequenced and assembled are listed in Table 1. *Lens ervoides* IG 72815 is the same accession used to develop the original reference genome, Ler.1DRT (Ramsay et al. 2021). CDC Greenstar is a large green lentil cultivar from the Crop Development Centre (CDC), University of Saskatchewan. All IG accessions were obtained from the genebank at the International Center for Agriculture Research in the Dry Areas (ICARDA) and have been maintained in the CDC collection for many years. BGE 016880 was the *Lens orientalis* parent of an interspecific RIL population generated by Fratini et al., 2007 and shared with the CDC. ILWL 25 is a *Lens nigricans* accession originally from ICARDA sourced from the Australian Grains Genebank (AGG).

**Table 1.** Species and accessions used for high molecular DNA extraction and genome assembly. Each genome code under “Genome Name” corresponds to the genome assembly version of each species.

| *Species* | *Accession* | *Material Source* | *Genome Name* |
| --- | --- | --- | --- |
| *Lens culinaris* | CDC Greenstar | CDC<sup>1</sup> | Lcu.1GRN |
| *Lens orientalis* | BGE 016880 | Fratini et al., 2007 | Lor.1WPS |
| *Lens odemensis* | IG 72623 | ICARDA<sup>2</sup> | Lod.1TUR |
| *Lens tomentosus* | IG 72805 | ICARDA | Lto.1BIG |
| *Lens lamottei* | IG 110813 | ICARDA | Lla.1ESP |
| *Lens ervoides* | IG 72815 | ICARDA | Ler.1DRT |
| *Lens nigricans* | ILWL 25 | AGG<sup>3</sup> | Lni.1VIC |

<sup>1</sup> Crop Development Centre at the University of Saskatchewan.
<sup>2</sup> International Center for Agriculture Research in the Dry Areas.
<sup>3</sup> Australian Grains Genebank.

DNA for all assemblies was derived from a single plant, or progeny of a single plant, that had been selfed for several generations. Plants were grown in controlled environment chambers, and leaves were harvested. For samples sequenced on the Gridion (Oxford Nanopore Technology), high molecular weight (HMW) DNA was extracted using a Salting Out Method (10x Genomics® Sample Preparation Demonstrated Protocol - Rev A, adapted from Miller et al., 1988) from preparations of intact nuclei (Zhang et al., 1995). For all samples used for Hi-C and PacBio Hi-Fi sequencing, HMW DNA was prepared using NucleoBond® HMW DNA kit (Macherey-Nagel, Düren, Germany). Genomic DNA for paired-end Illumina sequencing was extracted using E.Z.N.A.® Plant DNA DS Kits (Omega).

### Sequencing and pseudomolecule assembly of reference genomes

*Lens culinaris* genome Lcu.1GRN was assembled from 36.7x coverage of PacBio HiFi reads >10kb with LJA (https://github.com/AntonBankevich/LJA). Resulting contigs were scaffolded with an optical map generated with the Bionano hybrid scaffolding pipeline (https://bionano.com/) and then aligned to the *L. culinaris* Lcu.2RBY (Ramsay et al., 2021) assembly using mummer 3.23 (Kurtz et al., 2004) to create the pseudomolecules using a reference-guided method.

Sequencing platforms used and final depth of coverage for the wild genomes are described in Supplementary Table 1 and at https://knowpulse.usask.ca/research-study/wild-Lens-genome-assemblies. Assembly of Ler.1DRT was already reported in Ramsay et al. (2021). Assemblies of all other wild genomes were performed using the methodology fully described in the GenomeFocus workflow (https://github.com/ramsayl/genomefocus). Due to the lack of a genetic linkage map for *L. nigricans*, contigs were initially assigned pseudomolecule positions based on a reference-guided assembly using RagTag (Alonge et al., 2022) against the *L. ervoides* Ler.1DRT assembly (Ramsay et al., 2021), then manually corrected using the Hi-C data in Juicebox (Supplementary Table 1). Pseudomolecules were assigned chromosome numbers based on synteny with the *L. culinaris* chromosomes.

Minimap2 (Li, 2018) was used to align the genomes for subsequent synteny analyses. Major rearrangements in genome assembly order were confirmed at the chromosome level using oligo-FISH (Silvestrini et al., 2025).

### Repeat Analysis

*De novo* repeat annotation was performed using the EDTA pipeline (v1.9.4) (Ou et al., 2019), which identifies various classes of transposable elements (TEs), including long terminal repeat retrotransposons (LTR-RTs) and DNA transposons. The annotated TE sequences were classified using TEsorter (Zhang et al., 2022) to provide clade-level classification by searching against the REXdb database (Neumann et al., 2019). The resulting repeat library was then used to mask the genome with RepeatMasker (v4.1.6) (Tarailo-Graovac et al., 2009) to estimate the composition and distribution of repetitive elements across the genome assembly. Total satellite sequence estimation was performed using TideCluster (Novak, 2025) using default parameters.

### Gene annotation

Gene annotation was performed using the BRAKER2 pipeline, combining both *ab initio* and evidence-based methods. Protein sequences from *L. culinaris* and *L. ervoides* (Ramsay et al. 2021), were used as supporting evidence to annotate protein-coding genes on the soft-masked genomes. PASA (v2.5.3) (Haas et al., 2003) was then used to refine the gene models by incorporating transcript alignment evidence. Functional annotation of the predicted genes was carried out using BLAST+ (Camacho et al., 2009) and InterProScan (Jones et al., 2014) search against UniProt (The UniProt Consortium, 2023) and Pfam (Mistry et al., 2021) databases.

### Divergence analysis

Ks values were estimated for paralogous gene pairs to assess mutation rates and orthologous gene pairs between *Lens* species to estimate divergence times using Dated (version 1.0; https://github.com/ChuShin/dated), a Python-based script that employs the maximum likelihood method implemented in the PAML package (Yang, 2007). Orthologous genes were identified through reciprocal best BLAST analysis among single-copy colinear genes.

Divergence analysis was performed following the methodology outlined by Kagale et al., 2014. Mutation rates for each *Lens* species were estimated based on the average Ks (synonymous substitution) values associated with a shared whole-genome duplication (WGD) event, dated to approximately 56.5 million years ago. The mean mutation rate (8.33 × 10⁻⁹) across *Lens* species was then used to calculate divergence times between species (Supplementary Table 2).

To identify major peaks in Ks distributions, a Gaussian mixture model analysis was performed as described by Kagale et al. (2014). Briefly, Ks datasets were filtered to exclude values below 0.001, and histograms of log-transformed (ln) Ks values were generated with a bin width of 0.1. The R package Mclust (Scrucca et al., 2016) was used to fit Gaussian mixture models, from which the number of components (G), their means, and data proportions were derived. Skewed peaks in the Ks distributions, likely due to overlapping components, were resolved by merging overlapping Gaussians and representing them by their mean values. Divergence times between *Lens* species were estimated the mutation rate of 8.33 × 10⁻⁹ and the geometric mean Ks value of each distribution peak.

### Synteny analysis and rearrangement mapping

Synteny analysis was done with MCScanX (Wang et al., 2012). Rearrangement coordinate mapping between genomes was done using MCScanX output, considering protein sequences of annotated genes and excluding sequences associated with repetitive elements. Only BLAST hits involving primary transcript gene models were retained, while hits involving non-primary transcripts were removed to maintain consistency. The resulting MCScanX collinearity file was filtered using a custom Python script available at <https://github.com/mcbbaker/genome_synteny>. Different score thresholds were applied depending on whether blocks were intra-chromosomal (threshold: 1000) or inter-chromosomal (threshold: 5000) to selectively retain high-confidence syntenic blocks.

Due to residual noise and biological inconsistencies (e.g., gaps or overlapping blocks without known duplication events), manual curation was performed to remove spurious blocks and reintroduce a limited number of biologically plausible blocks from the original output. The final set includes some lower-scoring blocks, chosen to improve overall genome alignment fidelity between species. Results were visualized using Synvisio (https://synvisio.github.io).

Chromosomal rearrangement rates were calculated based on the ratio of the number of rearrangements per chromosome obtained from the rearrangement coordinates mapping, including inversions, translocations and inverted translocations, divided by species divergence time in million years as proposed by Nascimento & Pedrosa-Harand, 2023.

### Methylation calls

Oxford Nanopore Technology (ONT) raw reads for all genomes except Lcu.1GRN were aligned to the corresponding genome assembly using minimap2 (Li, 2018) (-x map-ont parameter). Nanopolish (Simpson et al., 2017) was then used to identify methylated (5mC) and unmethylated cytosines in the CpG context. CDC Greenstar PacBio reads were aligned against the Lcu.1GRN genome assembly using pbmm2 (https://github.com/PacificBiosciences/pbmm2) (--sort -j 100). Pb-CpG-tools (https://github.com/PacificBiosciences/pb-CpG-tools) was used for extracting methylation (5mC) calls in the CpG context from the aligned reads.

### Genome size estimates: Flow cytometry and KMer analysis

Young leaves of accessions listed in Table 1, except *L. nigricans* due to the lack of available seeds, were harvested at the size of 1 square centimetre and chopped using a razor blade in 100 μL of Extraction Buffer (CyStain UV precise P automate nuclei extraction and DNA staining kit, Sysmex, Kobe, Japan) to create a nuclei suspension. Young leaves of the endogenous control, *Raphanus sativus* (1.11pg/2C, Dolezel et al. 1992), were harvested at the size of 10 square centimetres and chopped using a razor blade in 3 mL of Extraction Buffer, with an additional 3 mL added after chopping to create a nuclei suspension. Each sample was then filtered using a CellTrics™ (Sysmex, Kobe, Japan) filter with 30 mm nylon mesh. For DNA size measurements, the control samples were mixed with 150 μL of DAPI staining solution, 20 μL of endogenous control nuclei suspension and 80 μL of sample nuclei suspension. 50 μL of nuclei suspension was added to 150 μL of DAPI staining solution. The measurements were done using a CytoFlex cytometer (Beckman Coulter, Brea - CA). Each sample size estimate was the mean of three repetitions analyzed on two different days.

Illumina short-read sequences were also used to estimate genome size. Quality-filtered reads (∼40× coverage) were processed with Jellyfish v2.2.7 using the parameters -C -m 21 -s 5G --min-quality=25 to generate a 21-mer frequency distribution. The resulting k-mer histograms were analyzed with GenomeScope (Vurture et al., 2017) to estimate genome size, heterozygosity, error rate, and repeat content.

To assess the relationship between genome size expansion and transposable element (TE) content across six *Lens* species, a Pearson correlation analyses were performed using R the package ggpubr (https://github.com/kassambara/ggpubr). Variables analyzed included total TE content, *Ty3-gypsy*, and *Ty1-copia* subfamilies, alongside genome size estimates obtained via flow cytometry.

### Immunofluorescence Assay

Seeds were germinated in the dark and harvested for nuclei and chromosome slide preparation, as in Oliveira et al., 2024. The roots were fixed at 10 °C in 3% Formaldehyde diluted in Tris buffer (10 mM Tris, 10 mM Na2EDTA, 100 mM NaCl) for 30 minutes with the first 5 minutes under vacuum and then washed in Tris buffer for 30 minutes on ice. The fixed roots were then used to prepare nuclei and chromosome suspension. The tissue was ground in in 1 mL of LB01 buffer (15 mM of Tris, 2 mM of Na2EDTA, 80 mM of KCl, 20 mM of NaCl, 0.5 mM spermine, 15 mM mercaptoethanol and 0.1% Triton X-100) using an Ultra-turrax T8 homogenizer. As primary antibodies P45, a custom-raised rabbit antibody against CenH3 (Neumann et al., 2015) and anti-α-tubulin raised in mouse (Sigma-Aldrich, St. Louis, MO; catalog number T6199) were used in the concentration of 1:1000 and 1:100, respectively, in 1xPBS with 0.1% Tween solution. The slides were incubated with both antibodies at 4 °C overnight. After incubation, the slides were subjected to two washes in 1xPBS for 5 minutes each and one wash in 1xPBS with 0.1% Tween for another 5 minutes. For the secondary antibody detection an anti-rabbit antibody conjugated with Rhodamine-red X (1:500 dilution; Jackson ImmunoResearch, Suffolk, UK; catalog number 111–295-144) and an anti-mouse antibody conjugated with Alexa488 (1:500 dilution; Jackson ImmunoResearch; catalog number: 115–545-166) were applied in the slides and incubated in the dark at room temperature for at least 1 hour. Another round of washes with 1xPBS and 1xPBS with 0.1% Tween was performed and the slides were fixed in 4% Formaldehyde in 1xPBS for 10 min at room temperature. One more round of washes with 1xPBS and 1xPBS with 0.1% Tween was performed, and the slides were mounted and stained with DAPI. Images were captured using a Zeiss AxioImager.Z2 epifluorescent microscope equipped with an AxioCam 506 mono-color camera and with an Apotome2.0 device and processed using Adobe Photoshop 24.1.1.

### Chromatin Immunoprecipitation sequencing (ChIP-seq) and centromere characterization

Intact nuclei from fresh leaves of all the analyzed species (Table 1) were extracted following the protocol of Neumann et al., (2012) and used to optimize the chromatin digestion with Micrococcal nuclease to the size of 1-3 nucleosomes. Chromatin digestion occurred with Microcal Nuclease (New England Biolabs, Ipswich, MA, USA) at a concentration of 200 U/uL from 30 to 40 minutes. After digestion, the chromatin was divided into two fractions, one of which was used as a control (input DNA) and the other fraction was used to immunoprecipitate the DNA associated with the centromeric histone H3 using the CenH3-2 antibody P45 (chipped DNA). The standard amount of CenH3-2 antibody used for ChIP was 15 μg/mL. To increase the specificity of the immunoprecipitation, the amount of the antibody was decreased to 3 μg/mL in duplicated experiments with *L. tomentosus*, *L. lamottei*, *L. ervoides* and *L. nigricans*. *L. culinaris*, *L. orientalis* and *L. odemensis* did not have duplicates because there was not enough leaf tissue. Chipped and Input DNA was sequenced at Admera Health (South Plainfield, NJ, USA) on Illumina NovaSeq X Plus platform producing 150 nt paired-end reads.

The Illumina sequencing reads were pre-processed using Trimmomatic (Bolger et al., 2014) to remove adapters and low-quality reads and then used for genomic mapping against the respective genome assemblies. The mapping software used was Bowtie2 (Langmead & Salzberg, 2012), together with SAMtools (Danecek et al., 2021), to deliver the output as a BAM file. A custom Python script was used to analyze the mapping output ( https://github.com/kavonrtep/cenh3_chip_seq_pipeline). The script contains different strategies for finding the best fit for the results, such as bin sizes of 200 bp and 2000 bp and three peak-calling software: macS3 (Zhang et al., 2008), Epic2 (Stovner & Sætrom, 2019), and bamCompare (Ramírez et al., 2016). The plotting was done using both IGV (Integrative Genomic Viewer) (Robinson et al., 2023) and JBrowse2 (Cain et al., 2022). The validation of the ChIP-seq experiments and the centromere composition analyses for each species was done using the RepeatExplorer2 pipeline and the ChIP-seq mapper tool implemented on a Galaxy server as described in Protocol 4 in Novák et al., (2020). The satellite similarity analysis was done using MAFFT (Katoh & Standley, 2013) and Muscle (Edgar, 2004), and the identity was calculated using P-distance. Satellite mapping was done using TideCluster (https://github.com/kavonrtep/TideCluster) with the centromeric satellites as a custom library. Bedtools (Quinlan & Hall, 2010) was used for centromere synteny and to assess the composition of transposable elements and cytosine methylation levels, using the start and end positions of each centromere (Supplementary Table 11).

Genetic maps used for centromere analysis had been previously generated for LR-100 *L. culinaris* x *L. tomentosus* (Stonehouse & Bett; https://knowpulse.usask.ca/genetic-map/Lens-interspecific/2026-RS-LR-100-superlinkage). Figures were generated using ggplot2 (R package) (Wickham, 2016), JBrowse 2 (Cain et al., 2022), IGV (Robinson et al., 2023) and Synvisio (https://synvisio.github.io/).

## Results

### Genomes of *Lens* species

We sequenced and assembled one genome of each of the seven species in the genus *Lens* (Table 1). This included a new *L. culinaris* assembly of cv. CDC Greenstar (Lcu.1GRN). The genomes were assembled with scaffold N50 ranging from 368 Mb to 507 Mb, and 94 % to 99 % of those scaffolds were anchored into the expected seven pseudomolecules for each *Lens* spp. Genome completeness was assessed with BUSCO (Manni et al., 2021), with scores ranging from 96.7 % to 98.9 %. Gene annotation identified 39,758 to 54,913 high-confidence genes, depending on species (Supplementary Table 2). Pseudomolecules were numbered into chromosomes among different species based on the synteny analysis with the published *L. culinaris* genome assembly (Lcu.2RBY).

### Genome size expansion is driven by transposable element content

Flow cytometry results show that the biggest genome is *Lens tomentosus*, at approximately 4.36 Gb, while the smallest is *L. ervoides,* at 3.34 Gb (Supplementary Figures 1 and 2, Supplementary Table 3). *L. nigricans*, which could only be calculated using K-mer analysis, has an estimated size of 3.03 Gb. All assembled *Lens* genomes are made up of more than 80 % transposable elements (TE) (Figure 1A and Supplementary Table 4). Long Terminal Repeats (LTR-RTs) are the main components, making up more than 70 % of each genome. *Ty3-Gypsy* elements are present twice as much as *Ty1-Copia*, with Ogre sequences being the most prominent type of *Ty3-Gypsy*, followed by Tekay elements.

**Figure 1.**
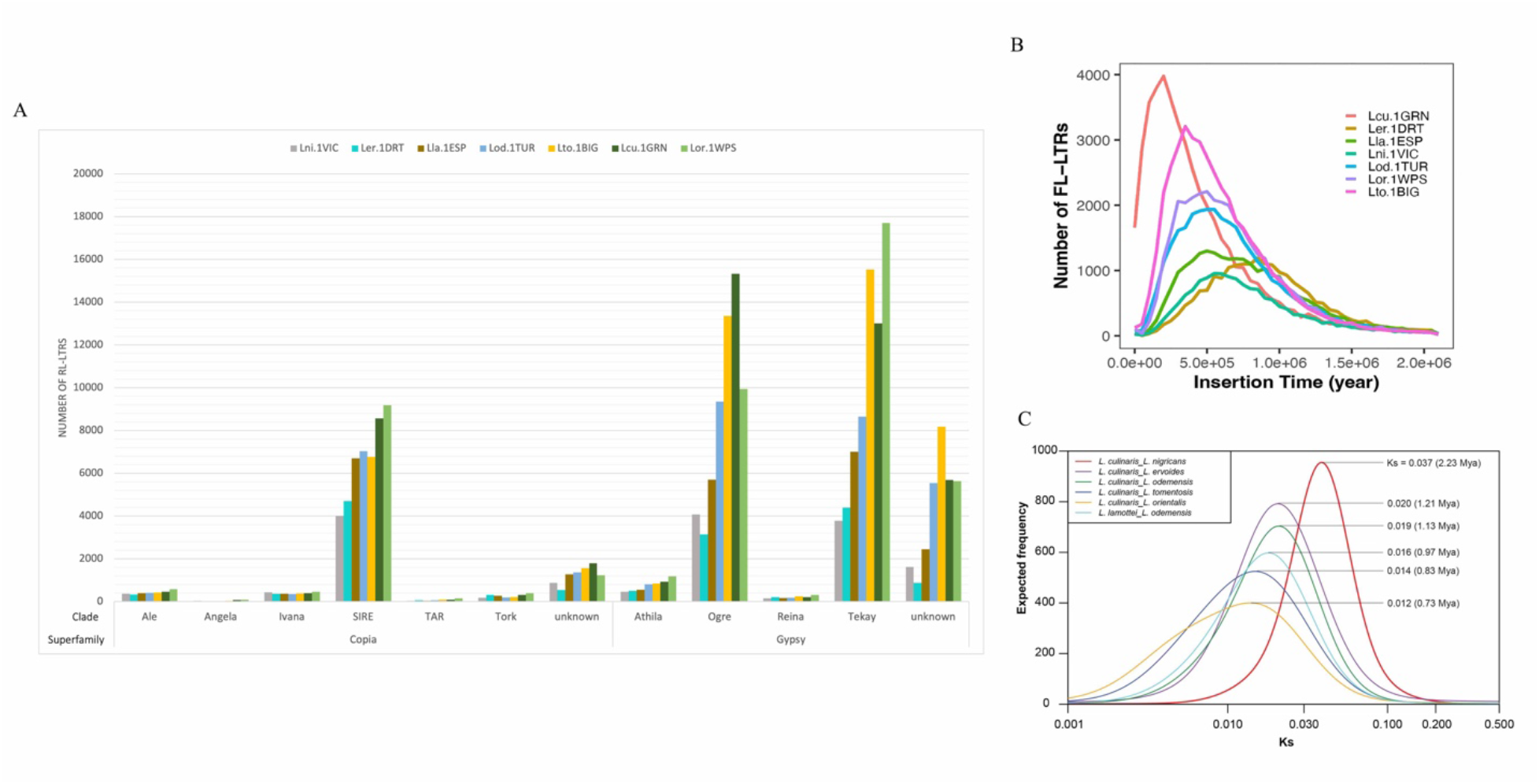
Transposable elements analysis and species divergence times in *Lens* genomes. A. Full-length LTR-RTs (Retrotransposons) composition of the most expanded Transposable Elements in *Lens* genomes. B. Transposable Elements (FL-LTRs) mean insertion times in *Lens* genomes in years. C. Estimated ages of divergence among *Lens* species in MYa.

Regression analyses of genome size variation with general and specific TE content were conducted to explore genome size expansion in *Lens* spp. (Supplementary Figure 3). There is a strong positive correlation between total TE content and genome size estimates, with these repeats accounting for more than 80 % of the genome size variation. *Ty3-Gypsy* alone explains nearly 65 % of the variation in genome size for all *Lens* genomes, led by Ogre and Tekay expansions (Supplementary Figure 3).

The TEs insertion times were determined for all genomes to identify TE burst patterns (Figure 1B and Supplementary Table 5). The most recent TE bursts are found in *L. culinaris*, while the oldest burst is present in *L. ervoides*; all other species fall in between, aligning with the phylogeny (Wong et al., 2015) and divergence (Figure 1C and Supplementary Table 6). These TE bursts correspond with the increased proportions of Ogre and Tekay (*Ty3-gypsy*) within *Lens* genomes, where recently differentiated species, such as *L. culinaris* and especially *L. tomentosus*, exhibit higher amounts of both TE clades compared to *L. ervoides* (Figure 1). *Ty1-Copia* content does not follow the same pattern as the TE bursts.

### Comparative centromere analysis shows diversity in *Lens* centromeres

Chromatin Immunoprecipitation Sequencing (ChIP-seq) experiments using P45 antibodies against the centromeric histone CenH3 (Robledillo et al., 2020) were done for all *Lens* species, allowing a comparative centromere analysis. P45 was initially developed to target *L. culinaris* CenH3 but BLASTP and the immunostaining assay confirmed the specificity of P45 against *L. culinaris* cv. CDC Greenstar and all wild *Lens* spp. genomes. At least one hit with 100 % homology was detected when using the P45 peptide against every *Lens* species peptide annotation (Supplementary Table 7).

The immunostaining assay confirmed the P45 antibody specificity and centromeric location for all *Lens* species, in which the mitotic α-tubulin, labelled in green in Figure 2, attaches to the CenH3 region, labelled in red with P45. The dot-like shape of the P45 signals matches the expected pattern for monocentric chromosomes for all *Lens* species (Figure 2).

**Figure 2.**
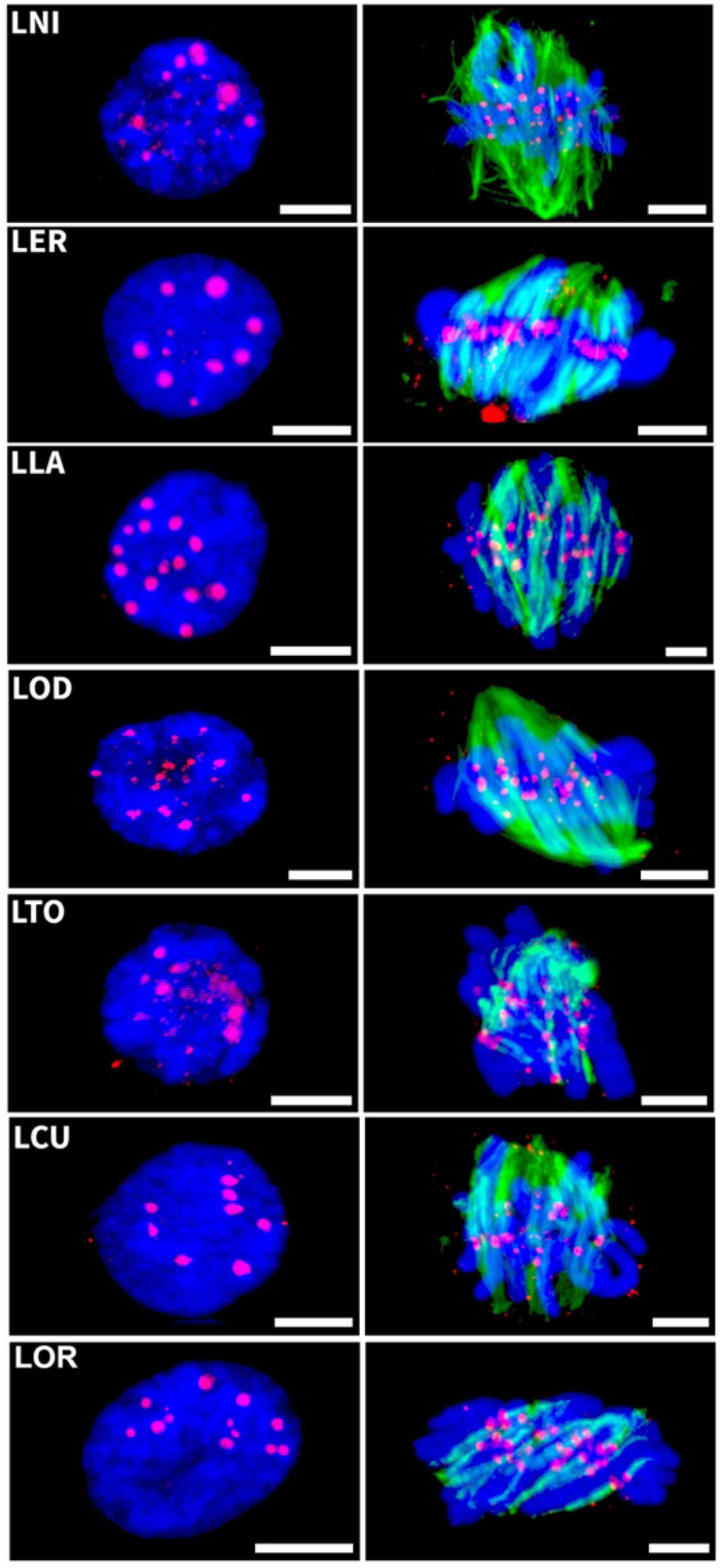
Immunostaining assay using P45 (anti-CenH3) labelled in red and anti-alpha-tubulin labelled in green. Chromatin is labelled with DAPI. LOR: *L. orientalis* BGE 016880, LCU: *L. culinaris* CDC Greenstar, LER: *L. ervoides* IG 72815, LLA: *L. lamottei* IG 110813, LNI: *L. nigricans* ILWL 125-0001, LOD: *L. odemensis* IG 72623, LTO: *L. tomentosus* IG 72805. First column shown nuclei signals; Second column shows mitotic metaphase signals. Bar: 10 uM.

There are 30 distinct centromeric satellite sequences spread across all the *Lens* chromosomes, with different proportions of each sequence in each species (Figure 3, Supplementary Table 8). Nine of the 30 centromeric satellites are shared across all species, and eight are species-specific. The more diverged species from *L. culinaris – Lens lamottei*, *Lens odemensis*, *L. ervoides* and *L. nigricans* (Figure 1C) tend to have higher numbers of species-specific satellites. This includes four of the five species-specific sequences, with only one species-specific satellite in *L. culinaris*. Species-specific satellites are less enriched than shared satellite families (Figure 3, Supplementary Table 8). Satellite sequences present in all seven genomes were likely present in the common ancestor of all *Lens* and remained conserved through *Lens* spp. differentiation.

**Figure 3.**
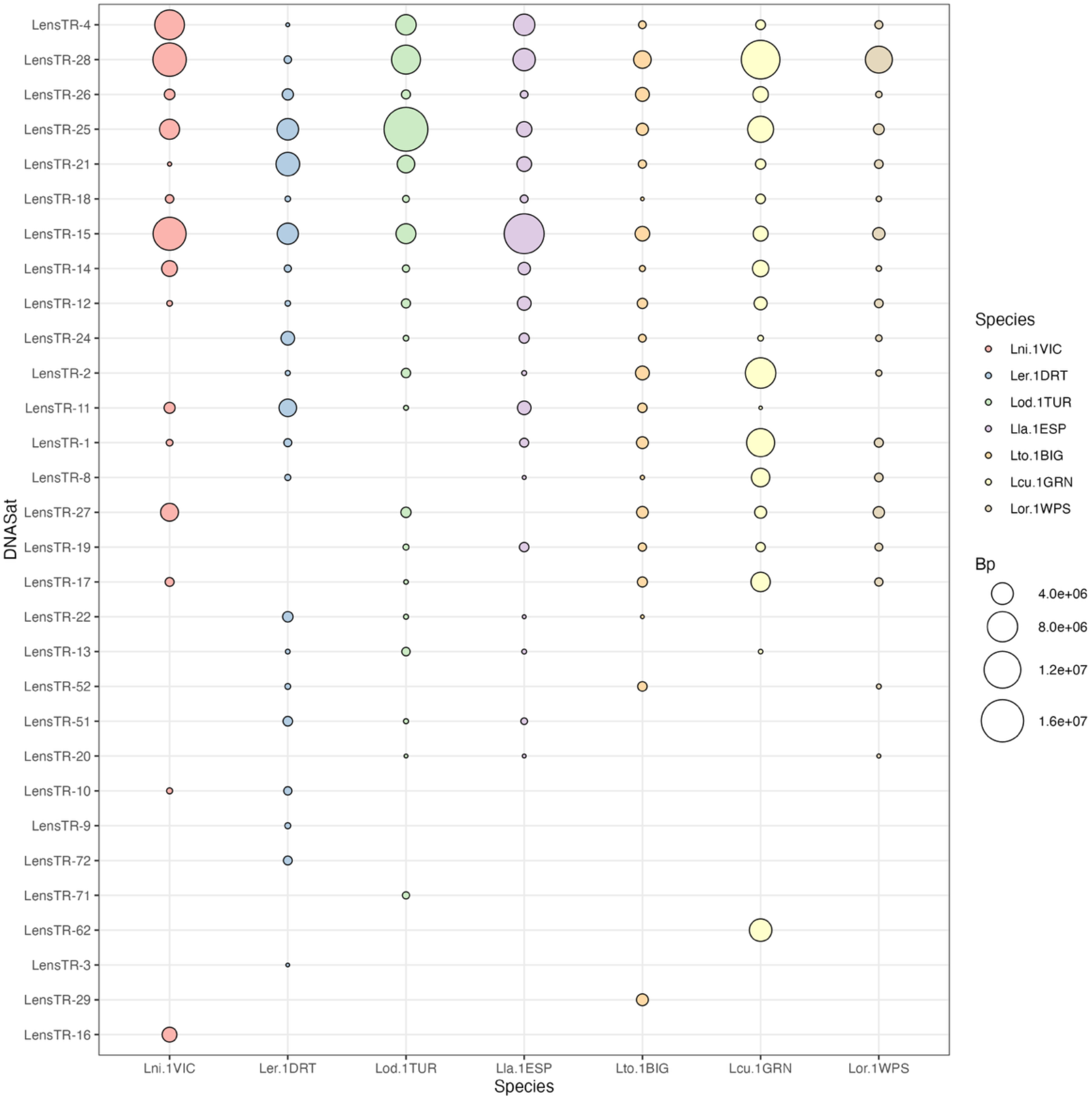
Presence and total base pair abundance of each centromeric satellite sequence across seven *Lens* spp. genomes based on ChIP-seq data analysis. Bp: The size of the circles represents the total base pairs of each satellite sequence in the whole genome data. Species codes according to Table 1.

On average, there are seven different centromeric satellite sequences in chromosomes 1 and 2 of all *Lens* and six different satellites in chromosomes 4, 5, 6, and 7. Chromosome 3 is the only one with over eight different satellite sequences (Supplementary Table 9). Four satellites were grouped into two different super-families (super-families 5 and 7) based on their higher homology to each other (over 80%). Super-family 7 is exclusively found in the secondary and tertiary *Lens* spp. genomes (Supplementary Table 8).

Satellites located exclusively within centromere coordinates are visualized in Figure 4 and differ from the whole-genome centromere satellite sequence composition in Figure 3, which can be distributed across different chromosomal regions. There is diversity even within a centromere of the same species, contradicting in part the centromere drive model, which requires a single centromeric satellite across all centromeres for a single species (Talbert & Henikoff, 2022). Some centromic satellites, such as LensTR-17 or LensTR-2, are conserved among closely related species. Other satellites are completely different, even within the same gene pool, such as LensTR-19 or LensTR-28, which are present in distinct centromeres of *L. orientalis*, *L. culinaris* and *L. tomentosus*.

**Figure 4.**
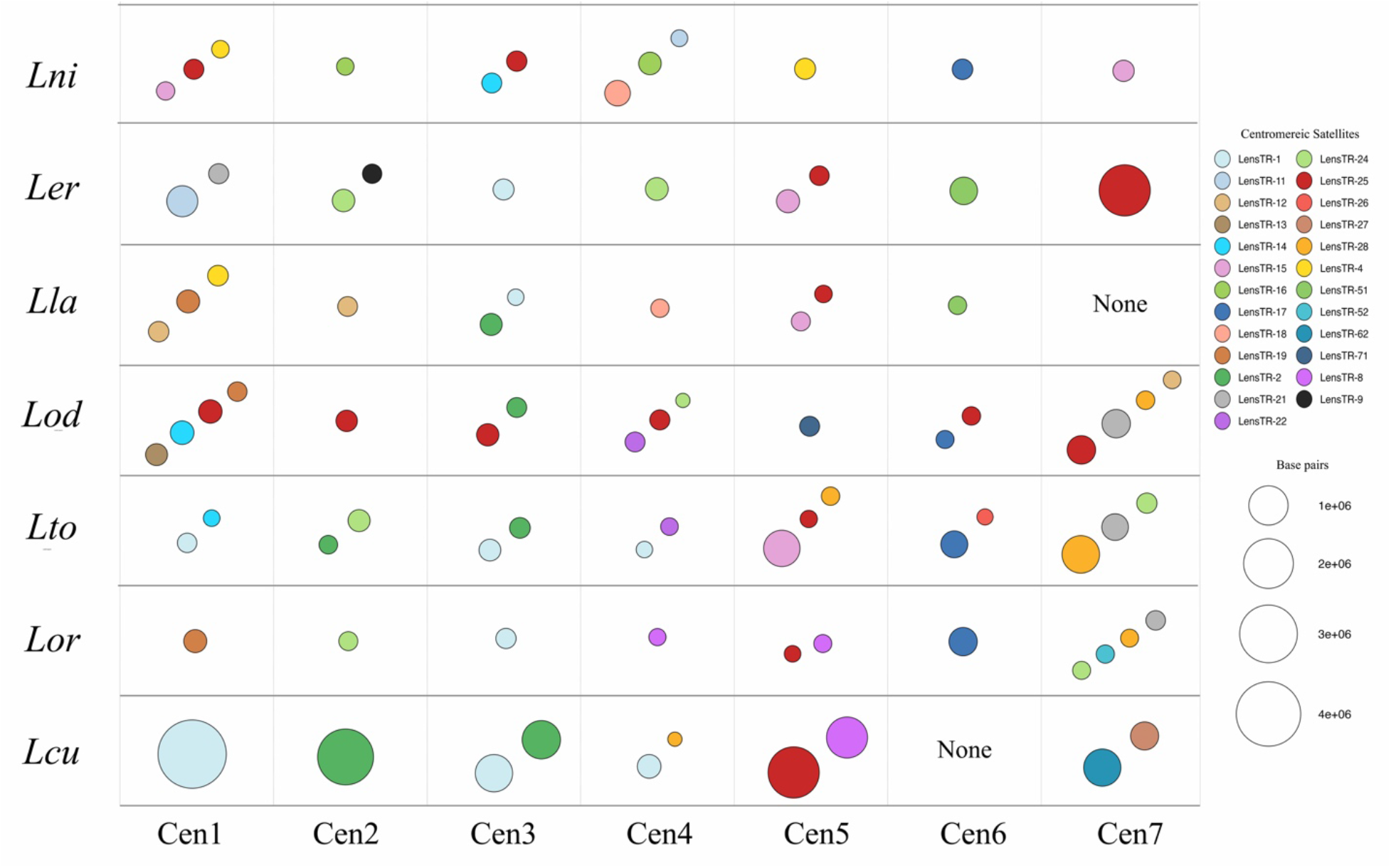
Centromere-located satellite sequence composition in *Lens* species by chromosome. Individual satellite names are coloured uniquely to show patterns of presence. None – no satellite sequence found. Size of circles represents total size of satellite in base pairs.

Legume species, including *Lens* spp., tend to have multiple centromeric satellite sequences within a species (Robledillo et al., 2020; Supplementary Figure 4). Those species also tend to have satellite sequences that are associated with centromeres (Figure 4) but are also located in regions outside the centromere (Figure 3) in other species. In total, all 30 centromeric satellite sequences identified in one or more *Lens* spp., mapped to at least one region outside the centromere in all *Lens* species, with 24 of those sequences being in at least one of the *Lens* spp. centromeres. The remaining six sequences are in unanchored contigs that are likely to be centromeric but are missing from the assemblies due to the challenge of assembling these highly repetitive regions (Supplementary Figure 5).

Sequence alignment of the *Lens* centromeric satellites with 64 previously published Fabeae centromeric satellites revealed higher levels of identity (> 90 %) with three centromeric satellites. Lens-TR8 had 99.27 % similarity with FabTR-35-LNS, LensTR-51 had 97.47 % similarity with FabTR-38-LNS and LensTR-71 had 93.83 % similarity with FabTR-36-LNS, classifying those satellites as the same centromeric sequences previously described for *L. culinaris* (Robledillo et al., 2020). None of the remaining satellites from the 13 Fabeae species had significant hits against the *Lens* satellite sequences.

All seven *Lens* spp. have hypermethylated centromeres, with cytosine methylation levels in CpG context (5mC) over 90 % within those regions (Figure 5A). *L. culinaris* had the smallest percentage (93.8 %) and *L. lamottei* the highest, with almost 99 % of its centromeric regions hypermethylated. *Lens* spp. centromeres are also enriched in LTR-RTs. On average, *L. orientalis* and *L. tomentosus* have the highest TE percentage in their centromeres (> 25 %), while *L. nigricans* has the lowest (15.45 %) (Figure 5B).

**Figure 5.**
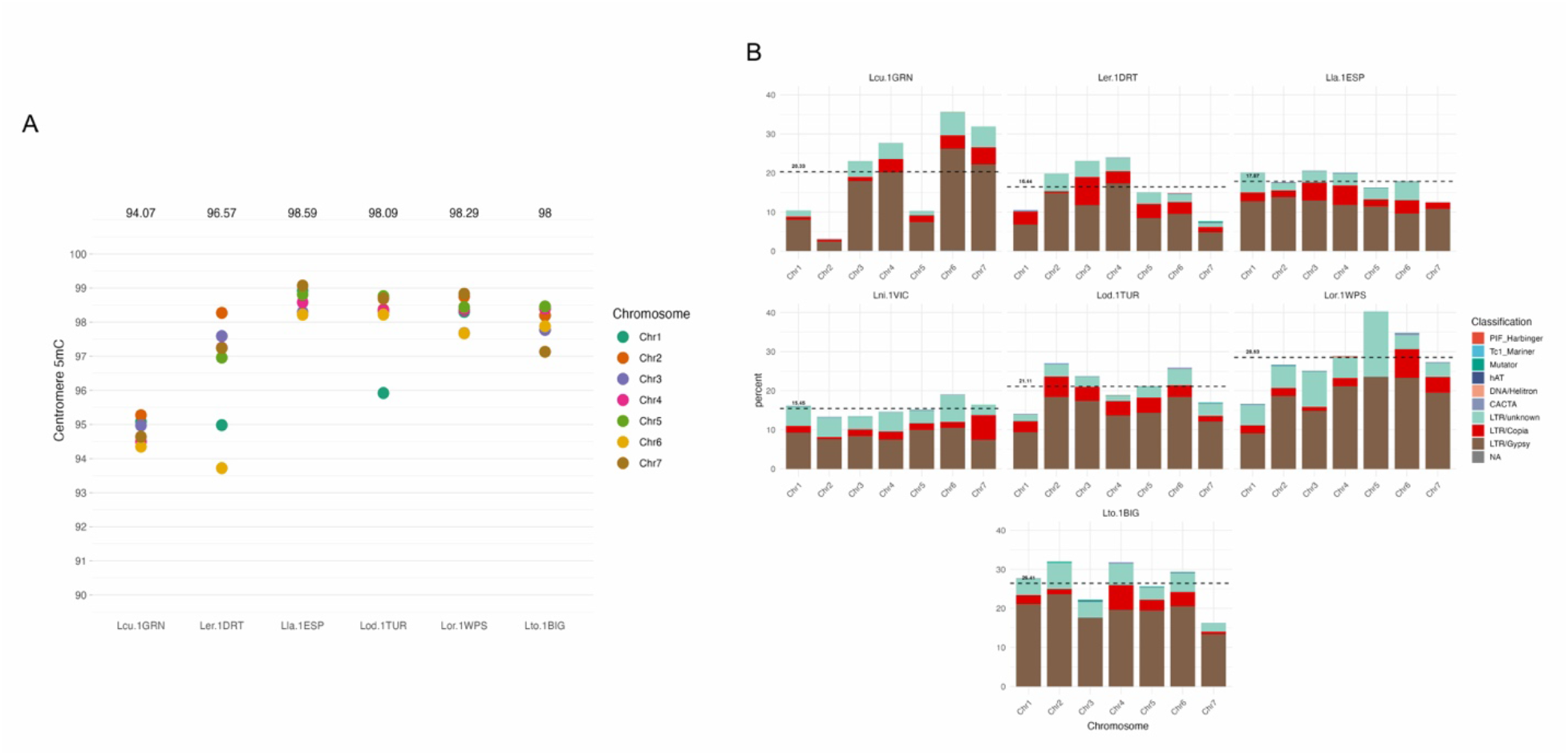
Cytosine methylation and transposable elements content for *Lens* centromeres by chromosomes. A. Centromere percentage of cytosine methylation (5mC) over C in CpG context across seven Lens genomes. Numbers above each genome represent species averages; B. Average transposable element (TE) content within *Lens* centromeres by chromosome. TE average across the seven chromosomes of each species is represented by dashed black lines.

### Synteny reveals evolutionarily fragile chromosomes and pericentric inversions events in *Lens* genomes

Synteny analysis among these genomes using homologous gene order reveals that many genomic rearrangements have occurred throughout *Lens* spp. evolution (Figure 6 and Supplementary Figure 6). Some rearrangements happened only in a single lineage, such as the large reciprocal Lni2 (Lni.1VIC chromosome 2)-Lni3 (Lni.1VIC chromosome 3) translocation and a large inversion on Lni1, distinguishing *L. nigricans* from the rest of the species. There are regions that appear to move more readily than others. For instance, the region at the distal end of Chr5 in Ler.1DRT, Lod.1TUR, Lcu.1GRN and Lor.1WPS is found at the distal end of Lla7 (Lla.1ESP) and is part of a larger translocation that is found at the distal end of Lto1 (Lto.1BIG). The major translocation distinguishing Lcu.1GRN from Lor.1WPS involves chromosomes 2 and 7. The proximal end of Chr7 in Ler.1DRT, Lla.1ESP, and Lor.1WPS is found at the distal end of Chr7 in Lod.1TUR and Lcu.1GRN, and the proximal end of Lto.1BIG Chr7. Lto.1BIG has double the average number of rearrangements relative to the other species (6.72 rearrangements, Table 2). This has been confirmed through oligo-FISH and is not an artifact of assembly errors (Silvestrini et al., 2025).

**Figure 6.**
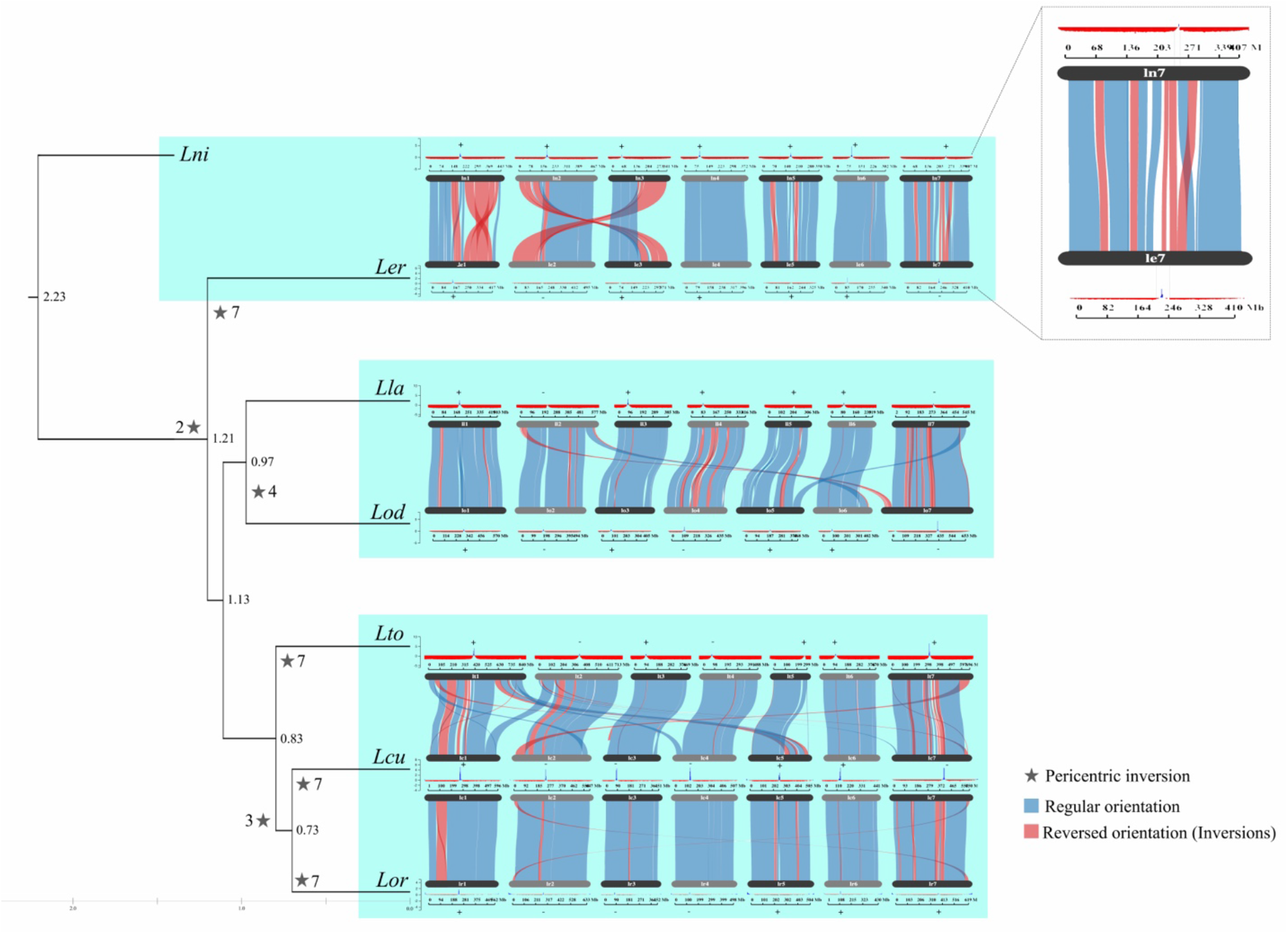
Gene synteny analysis reveals several rearrangements in *Lens* genomes. *Lens* species genomes are organized by phylogeny (Wong et al., 2015), in which the same genome appears more than once for centromere peaks visualization. Centromere peaks from ChIP-seq are shown on top of every *Lens* species’ chromosome. Inter-clade comparisons not included. Blue ribbons represent syntenic order of genes; red ribbons represent inverted order of genes in one chromosome relative to the other. Node values represent divergence time in MYa. Star numbers denote the chromosome involved in the centromere structural change. Lni and Ler chromosomes 7 are highlighted as an example of centromere peaks overlapping inverted regions. Signs of + and – above ChIP-seq peaks indicate structural rearrangement similarities in centromeres, whereas Lni is the reference with all centromeres classified as syntenic (+ sign).

**Table 2.**
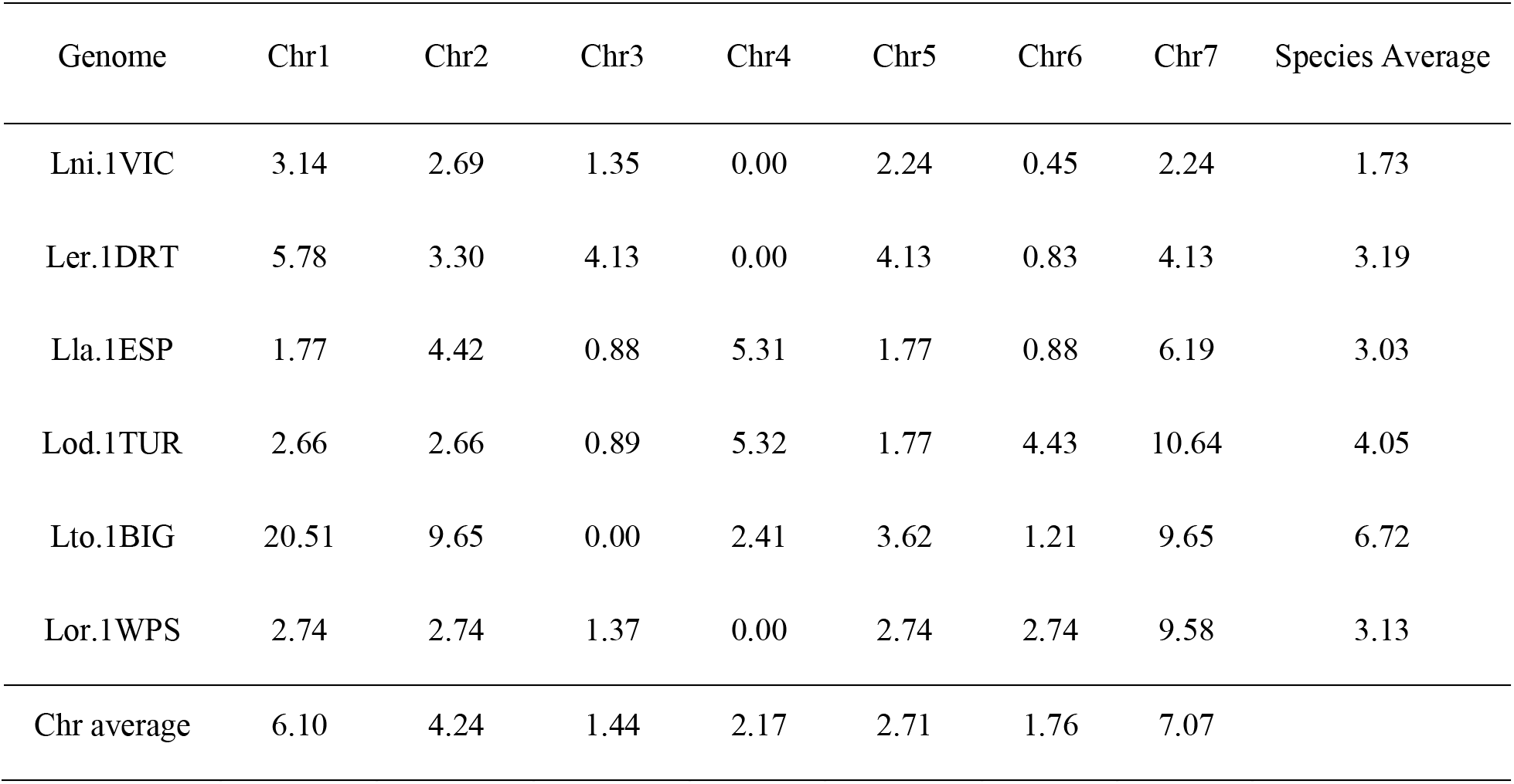
Chromosomal rearrangement rates per million years of six *Lens* species. Rearrangements include inversions, translocations and inverted translocations from Supplementary Table 10. Species divergence values from Figure 6.

| Genome | Chr1 | Chr2 | Chr3 | Chr4 | Chr5 | Chr6 | Chr7 | Species Average |
| --- | --- | --- | --- | --- | --- | --- | --- | --- |
| Lni.1VIC | 3.14 | 2.69 | 1.35 | 0.00 | 2.24 | 0.45 | 2.24 | 1.73 |
| Ler.1DRT | 5.78 | 3.30 | 4.13 | 0.00 | 4.13 | 0.83 | 4.13 | 3.19 |
| Lla.1ESP | 1.77 | 4.42 | 0.88 | 5.31 | 1.77 | 0.88 | 6.19 | 3.03 |
| Lod.1TUR | 2.66 | 2.66 | 0.89 | 5.32 | 1.77 | 4.43 | 10.64 | 4.05 |
| Lto.1BIG | 20.51 | 9.65 | 0.00 | 2.41 | 3.62 | 1.21 | 9.65 | 6.72 |
| Lor.1WPS | 2.74 | 2.74 | 1.37 | 0.00 | 2.74 | 2.74 | 9.58 | 3.13 |
| Chr average | 6.10 | 4.24 | 1.44 | 2.17 | 2.71 | 1.76 | 7.07 |  |

*Lens* spp. genomes seem prone to rearrangements, with specific chromosomes being particularly fragile and more often involved in those changes during *Lens* speciation. Among all seven chromosomes, Chr1, Chr2 and Chr7 are more fragile, while Chr3, Chr5 and Chr6 are the most stable throughout *Lens* evolution when looking at the chromosomal rearrangement rates per million years of divergence (Table 2).

Patterns among *Lens* chromosomes are also noticeable when analyzing genome proportions of cytosine methylation (5mC), transposable elements and total satellite sequence content (Supplementary Figure 7). For Lni.1VIC, Ler.1DRT, Lla.1ESP, Lod.1TUR, Lto.1BIG and Lor.1WPS genomes, chromosome 3 consistently presents the smallest proportions for all features, while chromosomes 2 and 7 exhibit the highest values.

ChIP-seq reads were mapped against the seven assembled genomes, producing the peaks shown in the tracks above the ribbons in Figure 6 and Supplementary Figure 6. Centromere coordinates are summarized in Supplementary Table 11. Each *Lens* species chromosome had at least one centromeric peak, with different enrichment values. Initially, Lcu.1GRN.Chr7 presented double independent CENH3 peaks. To confirm that it was a biological output, we analyzed the contigs in the specific centromere regions and detected an inverted contig, which will be further improved in the next release of this genome assembly. The remaining centromere analyses were based on the version containing the properly placed contig, thus resulting in a single ChIP-seq peak in chromosome 7 (Figure 6 and Supplementary Figure 6).

To compare the synteny of each centromere in relation to the most diverged species, *L. nigricans*, plus and minus signs above and below each ChIP-seq peaks (Figure 6) were included, where *L. nigricans* centromeres were treated as the ancestral state. Chromosomes 1, 5 and 6 were not involved in pericentric inversions; thus, all present the same orientation as *L. nigricans*.

Chromosome 2 differs in orientation between *L. nigricans* and the remaining *Lens* species, with the centromere located in the opposite orientation (minus sign), consistent with a pericentric inversion. The *L. culinaris*/*L. orientalis* clade had an inverted centromere on chromosome 3 compared to the remaining species. *L. odemensis* had an independent inversion in the centromere of chromosome 4; this was also evident in *L. tomentosus, L. culinaris* and *L. orientalis*. This could be a feature of the specific *L. odemensis* accession used in this study, as interspecific structural variation was already reported for *Lens* species (Silvestrini et al., 2025). Chromosome 7 had multiple independent pericentric inversion events, consistent with its designation as the most fragile chromosome in the genus (Table 2).

### Pericentric inversions are associated with segregation bias in *Lens* interspecies combinations

To test the effect of pericentric inversions on allele segregation in interspecific populations, a genetic linkage map of a Recombinant Inbred Line (RIL) population *L. culinaris* x *L. tomentosus* (LR-100; Stonehouse & Bett; https://knowpulse.usask.ca/genetic-map/Lens-interspecific/2026-RS-LR-100-superlinkage) was examined. Due to the presence of multiple translocations in *L. tomentosus* relative to *L. culinaris*, only linkage group (LG) 3 can be clearly separated, so it was chosen for allele bias analysis and centromere synteny analysis (Figure 7 and Supplementary Figure 8). *L. culinaris* centromere positions in chromosome 3 were compared to *L. tomentosus* centromere 3 according to the gene synteny order of each parent. A pericentric inversion was detected in the cultivated species when compared to *L. tomentosus*. Allele segregation shows a clear bias towards *L. tomentosus* in the pericentric inverted region of chromosome 3, different from the remaining sections of LG3 (Figure 7).

**Figure 7.**
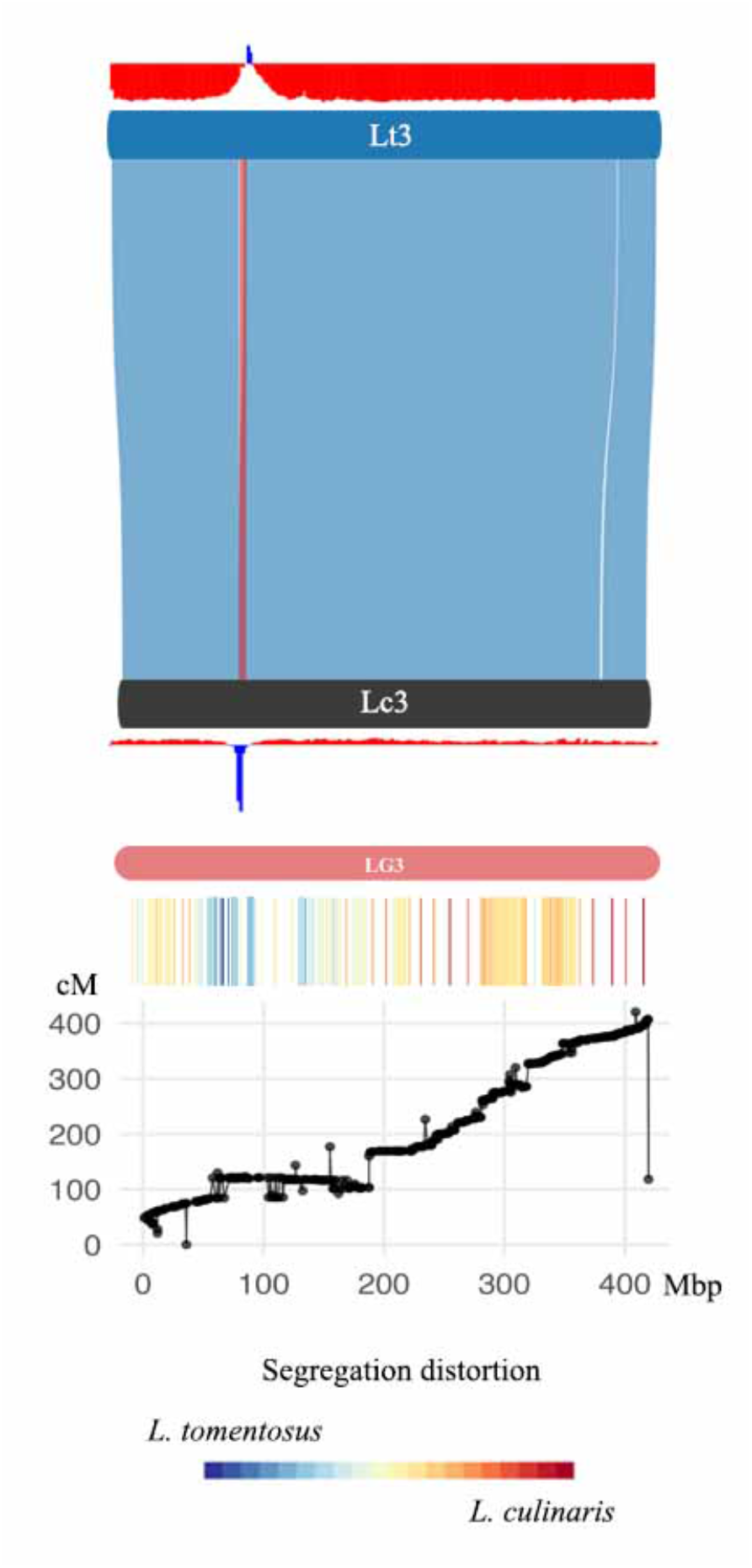
Linkage map analysis of a pericentric inversion detected in chromosome 3 in *Lens culinaris* and *Lens tomentosus* interspecific combination. First level shows synteny between wild parents (Lt) and cultivated (Lc). Second level shows the genetic map with linkage groups (LG). Third level shows allele frequency of one parental over the other. Fourth level shows the plotted recombination frequency vs. physical distance to determine the centromere position in lower recombining areas. The remaining linkage groups are found in Supplementary Figure 8. Chromosome 3 was chosen for plotting centromere positions due to the clear separation in one linkage group when compared to the remaining chromosomes. Blue ribbons indicate regular orientation of gene order, and red ribbons indicate inverted orientation of gene order.

## Discussion

We sequenced and assembled the genomes of at least one each of cultivated lentil and its six wild relatives. This is important considering the limited availability of whole-genome resources for *Lens* (Ramsay et al., 2021) and the importance of wild relatives in broadening the genetic diversity of cultivated lentil (Hajjar & Hodgkin, 2007, Coyne et al., 2020). The genus *Lens* also presents a good resource for evolutionary genomics studies due to the diploid nature and small number of chromosomes (n=7) of all members of the genus. An analysis of gene synteny across the genus showed that specific chromosomes—*viz*. those syntenic with Lcu 1, 2, and 7, are more fragile and prone to rearrangement than the others. Distinct patterns of repeats and 5-methylcytosine were detected among *Lens* spp. chromosomes, particularly in fragile chromosomes such as 2 and 7 and the most evolutionarily stable one, chromosome 3.

Although very similar, the genomes of humans and chimpanzees differ by inversions and translocations related to at least ten chromosomes that eventually led to our main speciation event (Suntsova & Buzdin, 2020). Navarro & Barton, 2003 propose that these ten rearranged chromosomes evolved at a different pace compared to fully syntenic chromosomes. Our results suggest a similar differential evolutionary pace for rearranged chromosomes in *Lens* genomes. Evolutionarily fragile chromosomes have also been reported in other organisms, such as insects and other plant species (Brown & O’Neill, 2010; Ruiz-Herrera & Robinson, 2007).

Of the ten chromosomes proposed to be involved in a faster evolutionary pace between humans and chimpanzees, nine of those chromosomes are also reported to be involved in pericentric inversions, either from one species or the other. Pericentric inversions are known to cause genomic instability, mainly due to alternative meiotic configurations during Prophase I, which can induce more rearrangements (Bhat & Wani, 2017; O’Neill et al., 2004; Rieseberg, 2001; Termolino et al., 2019). *Lens* spp. fragile chromosomes have pericentric inversions in at least five species. Those pericentric inversions could explain the fragility and the propensity to rearrange of some of the *Lens* spp. chromosomes.

Allele segregation data from an interspecific RIL population derived from a cross between a *L. culinaris* accession and a *L. tomentosus* accession are distorted towards the parent that does not have a pericentric inversion. Although previous studies have suggested that centromere drive is absent or occurs only episodically in *Fabeae* due to centromeric sequence diversity (Robledillo et al., 2020), the observed bias toward one parental allele could represent one such episodic event. Centromere drive can result in preferential segregation of alleles from one parental chromosome, especially around the centromere, when changes like pericentric inversions result in differences in centromere size in one of the parental centromeres (Huang & Rieseberg, 2020; Searle & de Villena, 2022). Lui et al., (2023) demonstrated that different centromeres in a parental combination of another legume species can result in differences in the centromere composition of the progeny, which likely explains the bias seen in our results for chromosome 3. Centromere drive can act to either fix or eliminate the chromosome with the pericentric inversion, as seen in other Eukaryotes (Finseth et al., 2021; Searle & Villena, 2022).

Legume species are known to exhibit diverse centromeres in terms of both structure and sequence composition (Macas et al., 2023; Neumann et al., 2015; Robledillo et al., 2020). Species belonging to the genera *Pisum* and *Lathyrus* possess elongated centromere structures referred to as metapolycentric chromosomes, while *L. culinaris* and other species within Vicia are reported to have monocentric chromosomes (Macas et al., 2023; Neumann et al., 2012, 2015). Our results include six new *Lens* species, all of which displayed dot-like CenH3 signals, further attesting to the monocentric nature of the *Lens*/*Vicia* clade. Robledillo et al., (2020) reported 64 satellite sequences associated with centromeric chromatin in 14 Fabeae species, most of which are noted to have multiple centromeric satellite sequences, even within the same centromere. Our results suggest similar findings for *Lens* species in which centromeres of *L. odemensis* possess up to four different satellite sequences each. The presence of satellites associated with centromeric chromatin in one species, and with non-centromeric chromatin in other species, as detected in *Lens* spp., has also been reported (Robledillo et al., 2020).

The source of diversity in centromeric satellite sequences in legumes has also been discussed, with several hypotheses being dismissed (Macas et al., 2023; Robledillo et al., 2020). The lack of genome structural comparisons among closely related legumes has led to the proposition that legumes have a “stable karyotype” and a disregard for the extent of structural variation in generating centromere diversity in those species (Robledillo et al., 2020). *Lens* spp. genomes are highly rearranged, with inversions including the centromere region, challenging this premise. Macas et al., (2023) detected structural variation within centromeres of seven Fabeae species, Liu et al., (2023) detected centromere repositioning in soybeans and Wei et al., (2025) detected inversions associated with centromere structure in Phaseoleae species. Those authors, in conjunction with this study, contribute to an enhanced understanding of legume centromere structural diversity.

Genomic rearrangements may explain the dynamism and sequence diversity of legume centromeres, promoting TE expansion (Raskina et al., 2008) and accelerating concerted satellite evolution, sometimes with a different “pace” between chromosomes of a same species as seen in our results, which leads to inter- and intra-species centromere satellite diversification (Hartley & O’Neill, 2019). We found evidence of differential segregation bias of a centromere involved in pericentric inversions, which might suggest the occurrence of centromere drive associated with chromosomal rearrangements rather than sequence composition itself in a legume species. With advances in legume structural and comparative genomics, new insights may provide a broader perspective on the role of chromosomal and centromere rearrangements in centromere satellite sequence diversity in this group of plants.

## Acknowledgments

This work was supported by the Enhancing the Value of Lentil Variation for Ecosystem Survival (EVOLVES) project funded by Genome Canada [grant: LSP18-16302] and managed by Genome Prairie. Matching funding was provided by: Western Grains Research Foundation [grant: GC1903], Saskatchewan Ministry of Agriculture [grant: 20200026], BASF, and the University of Saskatchewan. *L. orientalis* sequencing was funded in part by the Canada First Research Excellence Fund (CFREF) and Plant Phenotyping and Imaging Research Centre (P^2^IRC), managed by the Global Institute for Food Security (GIFS). *L. nigricans* work was funded by Agriculture Science and Technology, Melbourne, Victoria, Australia. We thank Dr. Jiri Macas’ group for the experimental molecular biology space and support from the Czech Science Foundation grant 24-10036S. Computational resources and data storage facilities were in part provided by the ELIXIR-CZ Research Infrastructure Project (LM2023055). EvW is supported by the Vermont Agricultural Experiment Station under USDA NIFA NE2210 and USDA ARS Food Systems Research Center.

## Data Availability

Genome assembly FASTA files are available via NCBI BioProject PRJNA1520023. Genome assembly FASTA files and JBrowse 2 tracks containing gene annotations, centromeric peak calls, 5mC methylation in the CpG context, repetitive sequence annotations, and genome synteny are also available through KnowPulse at https://knowpulse.usask.ca/experiment/EVOLVES-Lens-centromeres.

