## Supplementary Figures for "*Lens* species genome assemblies reveal evolutionarily fragile chromosomes associated with pericentric inversions"

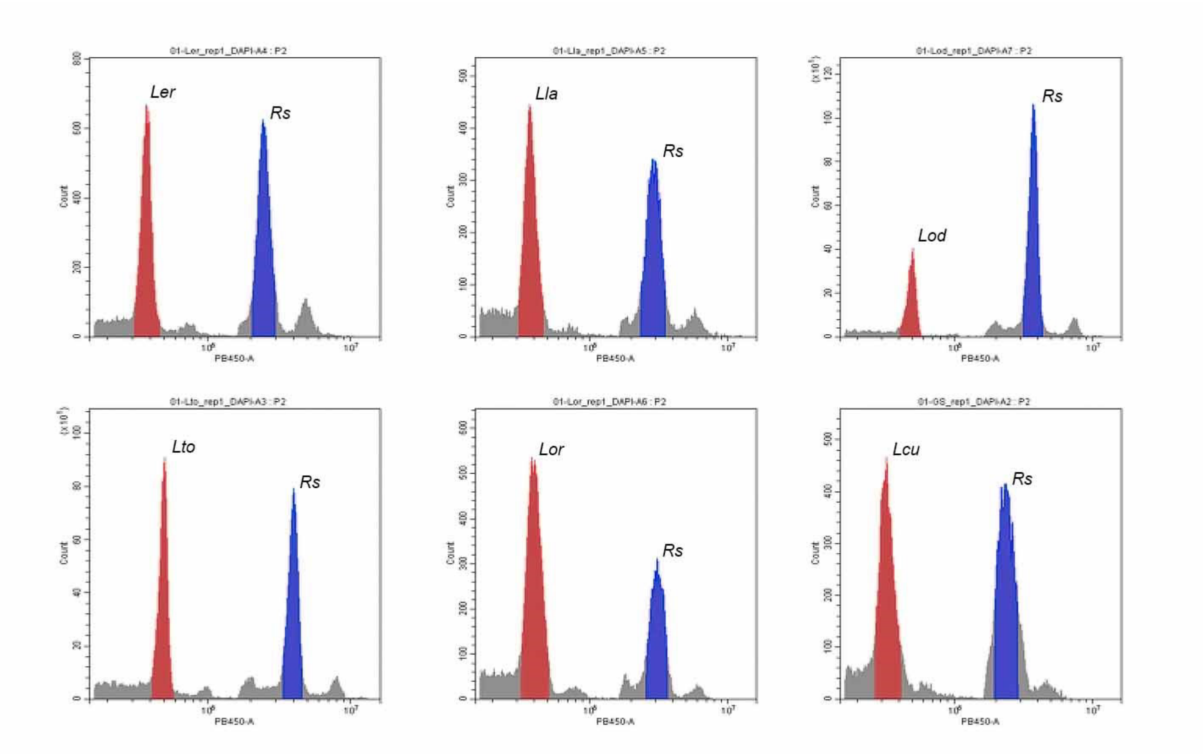


Supplementary Figure 1. Flow Cytometry 2C ploidy level peaks used to estimate the size of each *Lens* spp. Ler: *Lens ervoides*, Lla: *L. lamottei*, Lod: *L. odemensis,* Lto: *L. tomentosus*, Lor: *L. orientalis*, Lcu: *L. culinaris*. Rs: *Raphanus sativus* endogenous control with a genome size of 1.11pg/2C (Dolezel et al., 1992).

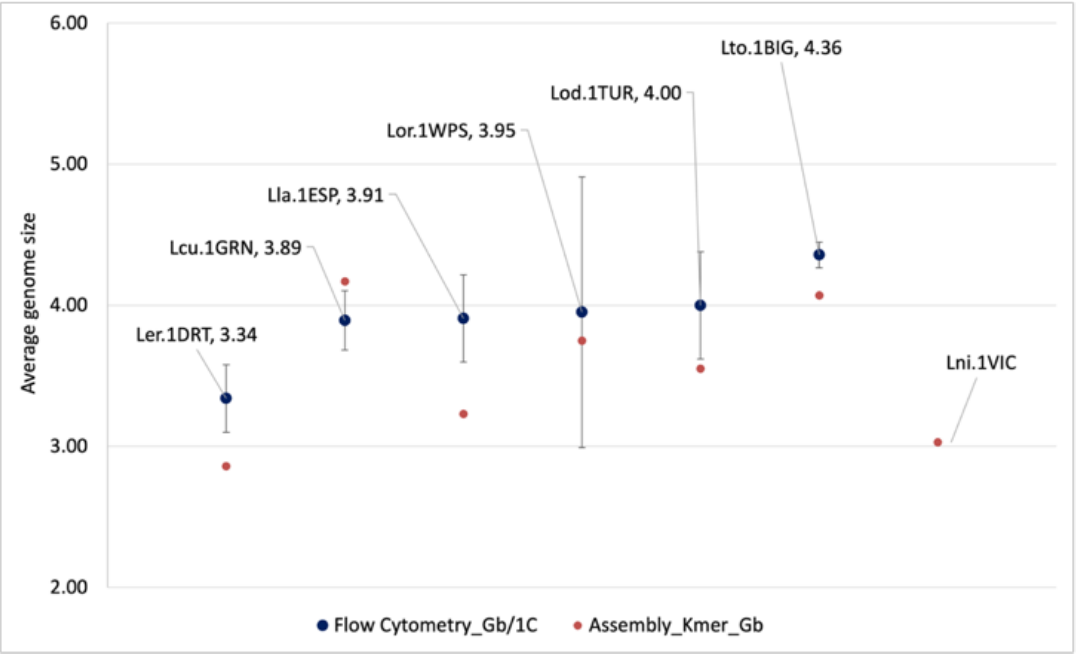


Supplementary Figure 2. Genome size estimation for *Lens* spp. genomes using Flow Cytometry and

Kmer Analysis. Bars represent the standard deviation from the mean of different measurements in the Flow Cytometry.

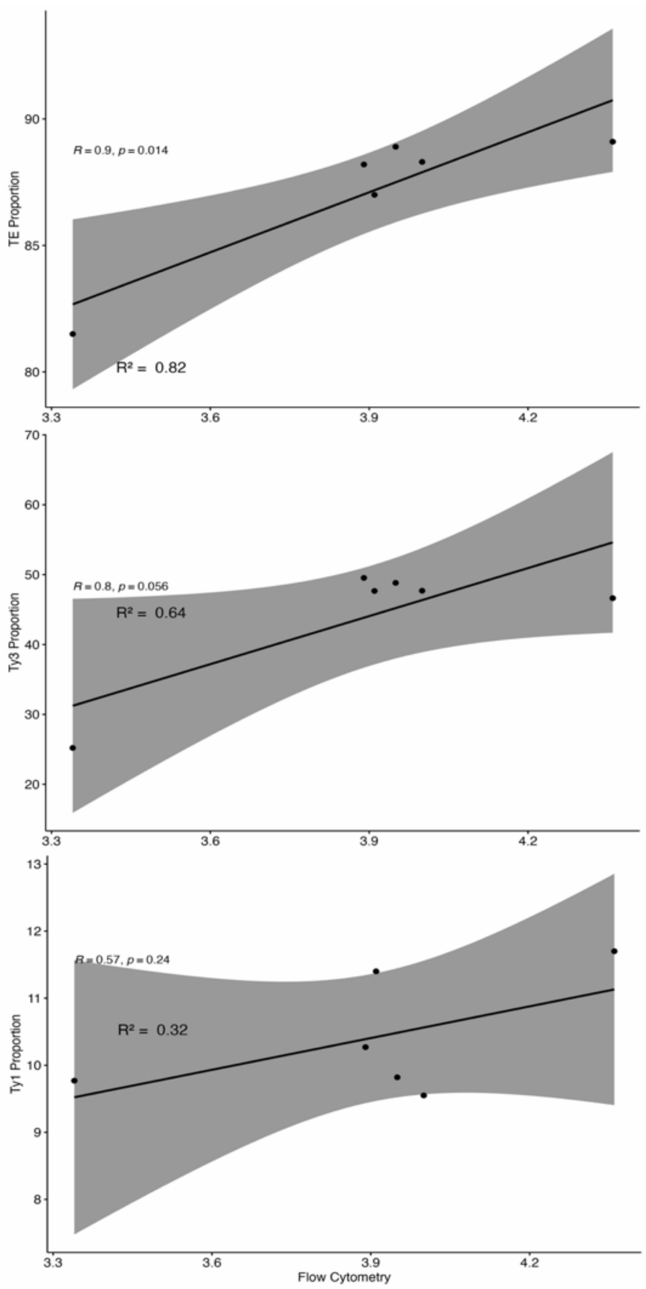


Supplementary Figure 3. Regression analyses of flow cytometry genome size variation and repeat content.

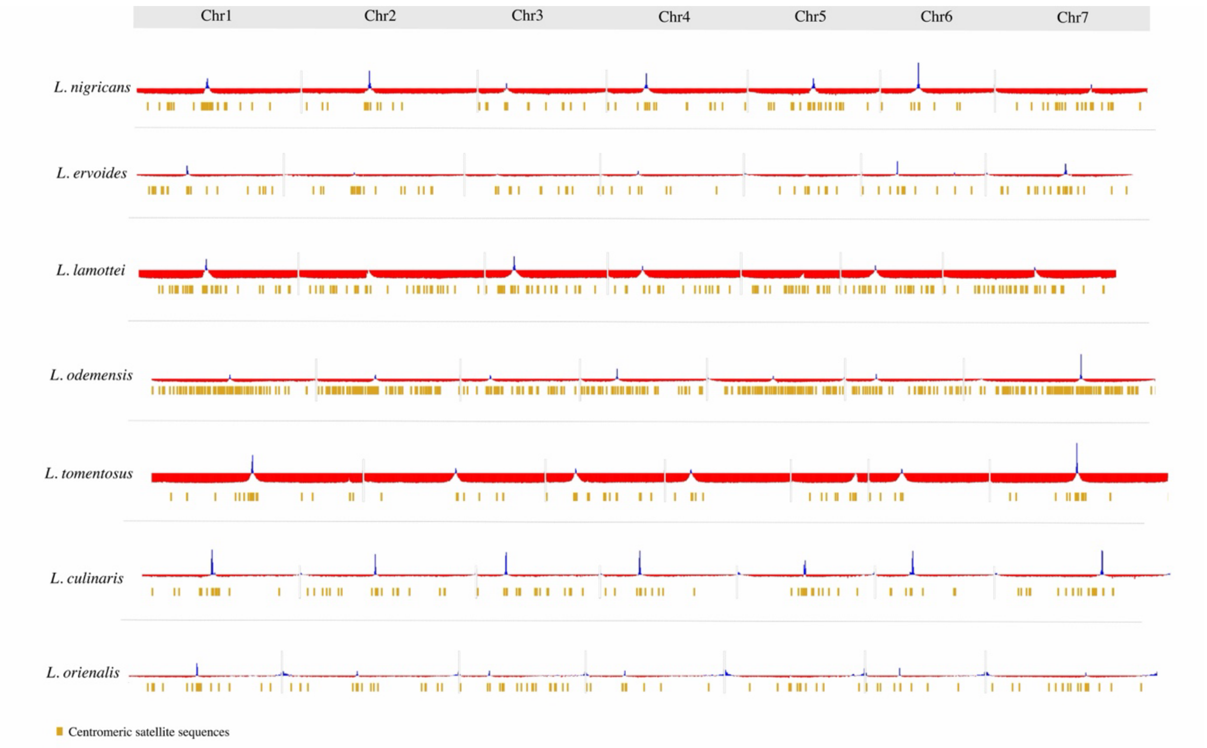


Supplementary Figure 4. Centromeric satellite sequences located outside the centromere region in the seven *Lens* species.

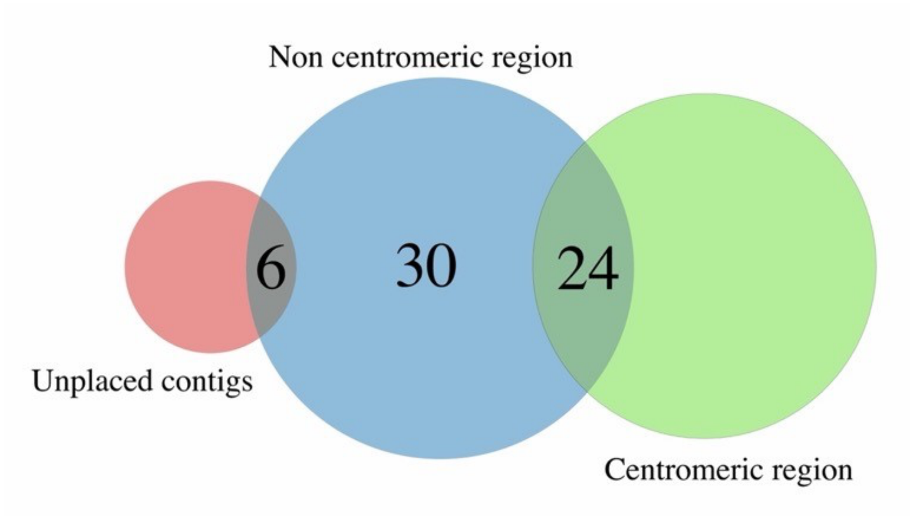


Supplementary Figure 5. Venn diagram of the number of satellite sequences that are centromeric in at least one species (green circle) and the number of satellites that are found outside the centromere (blue circle). Not all satellites were found in at least one centromere due to unplaced contigs (red circle).

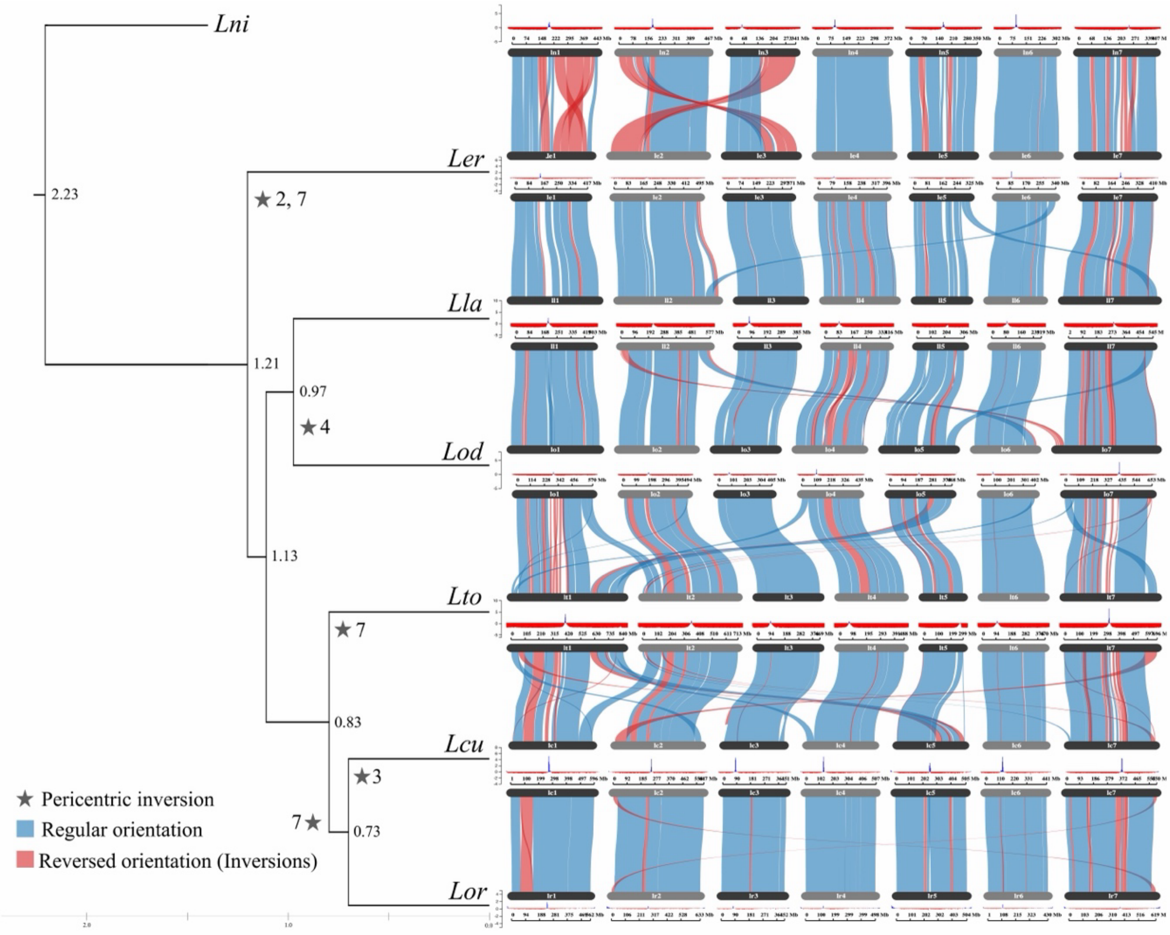


Supplementary Figure 6. Complete synteny and centromere comparisons among *Lens* species. Includes inter-clades comparisons.

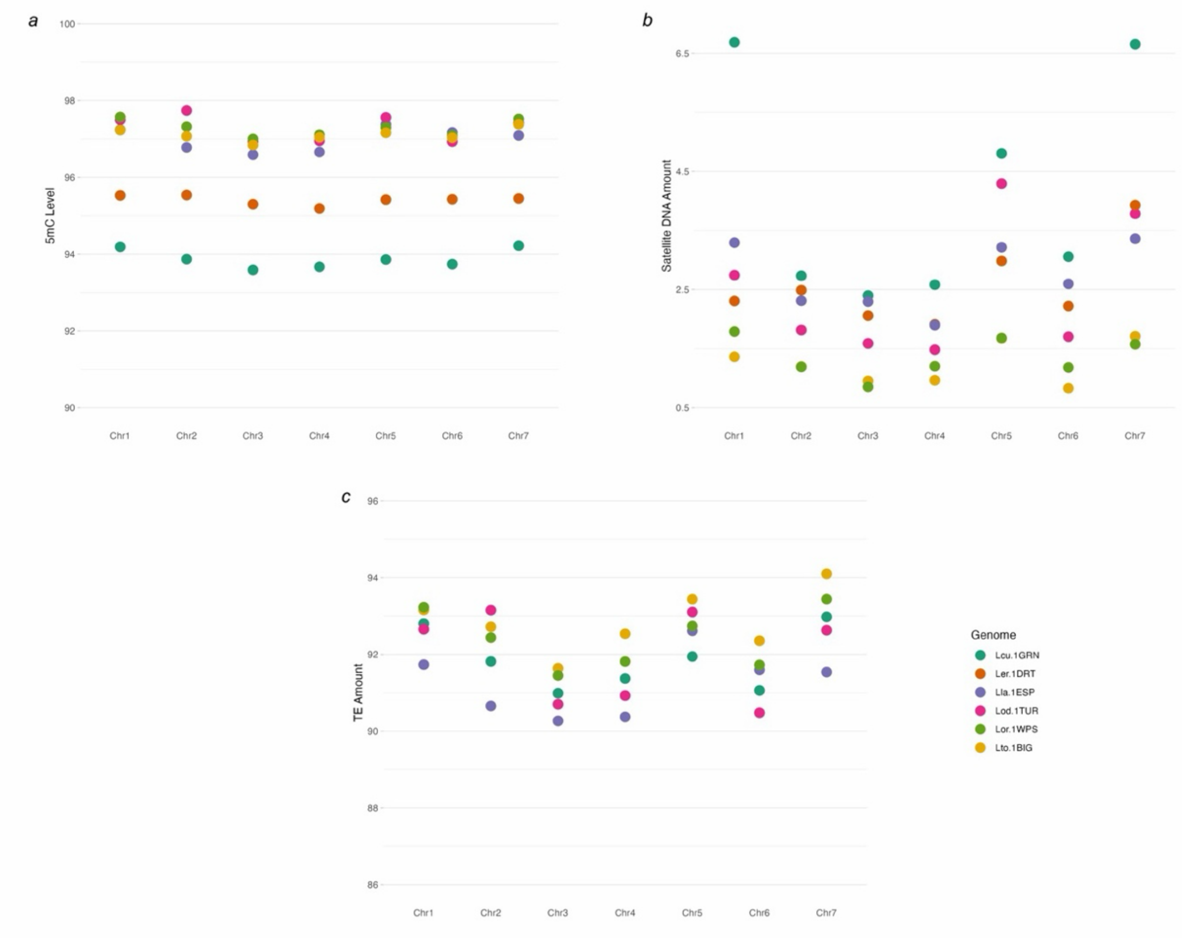


Supplementary Figure 7. Distribution of (A) Cytosine Methylation (5mC) in percentage of 5mC over C per chromosome, (B) Total satellite proportions in percentage related to each chromosome and (C) Transposable elements in percentage related to each chromosome across individual chromosomes in six *Lens* genomes.

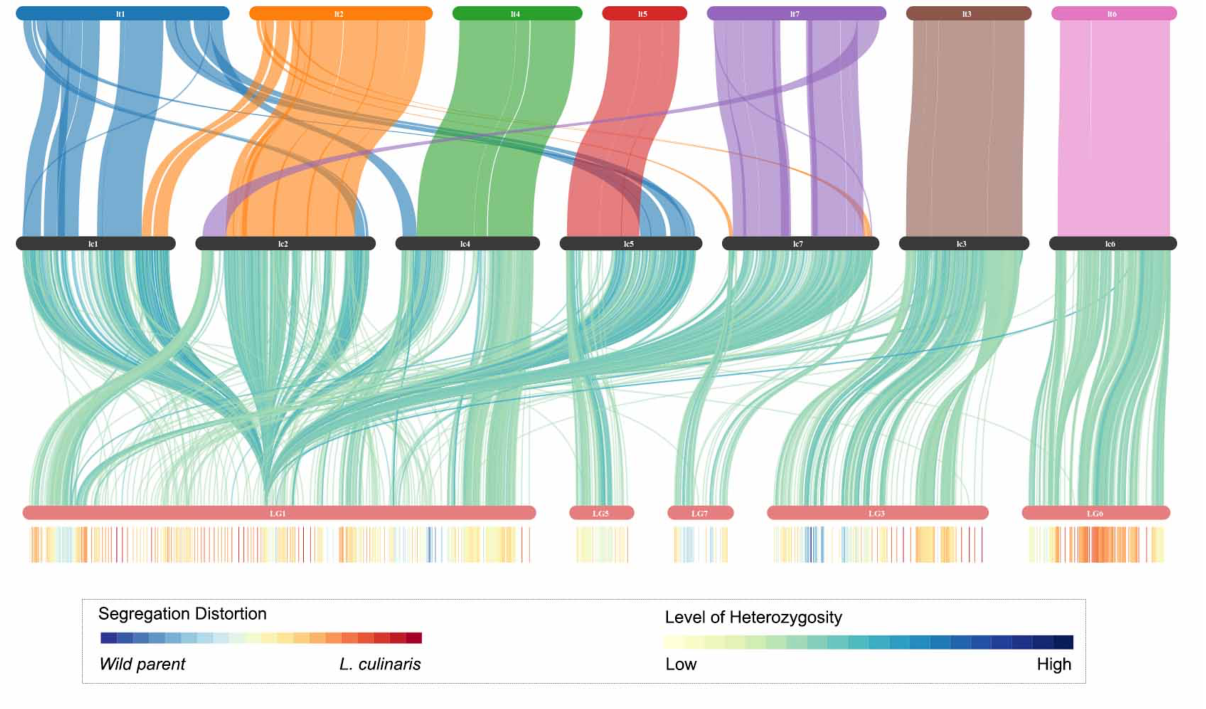


Supplementary Figure 8. Linkage map analysis in *Lens* interspecies combinations. First level shows synteny between wild parents (Lt) and cultivated (Lc). Second level shows the genetic map with linkage groups (LG). The third level shows heterozygosity levels and allele frequency of one parental over the other.
